# Monitoring tropical forests with light drones: ensuring spatial and temporal consistency in stereophotogrammetric products

**DOI:** 10.1101/2025.06.27.661894

**Authors:** Nicolas Barbier, Pierre Ploton, Hadrien Tulet, Gaëlle Viennois, Hugo Leblanc, Benoît Burban, Maxime Réjou-Méchain, Philippe Verley, James Ball, Denis Feurer, Grégoire Vincent

## Abstract

Light drones provide a cheap and effective tool to monitor forest canopy, especially in tropical and equatorial contexts, where infrastructure and resources are limiting. In these regions, good quality optical satellite images are rare, yet the stakes are maximal to characterize forest function, dynamics, diversity, and phenology, and more generally the vegetation-climate interplay.
We describe a complete processing chain based on photogrammetric tools that seeks to optimize the spatial and spectral coherence between repeat image mosaics at centimetric resolution. Our target is to allow individual tree-level monitoring over tens to hundreds of hectare scales with consumer grade equipment (i.e., quadcopter with stabilized RGB camera, standard GNSS positioning).
We demonstrate the increase in spatial precision achieved using Time-SIFT and Arosics algorithms, which allow (individually and synergistically) to reduce global and local spatial misalignment between mosaics from several meters to a few centimeters. Time-SIFT provides the advantage of increased robustness in initial image alignment and 3D reconstruction, and hence reduces occasional distortions or data gaps. Using Agisoft’s color and white balance corrections combined with the use of vegetation indices provides meaningful quantitative signal despite considerable changes in acquisition conditions.
In particular, indices that are less sensitive to illumination changes, like the green chromatic coordinate (GCC), allowed evidencing a seasonal signal over four years of monitoring in the evergreen moist forest at Paracou in French Guiana. The signal was decorrelated from obvious geometrical effect (sun height), and provided information on the vegetative stage at tree, species, and stand levels.

**Data/Code for peer review:** The complete processing chain, as well as the data and scripts used for producing the analyses presented here, are available for review on Zenodo: https://zenodo.org/records/15449377?token=eyJhbGciOiJIUzUxMiJ9.eyJpZCI6IjdjNWExMzIzLThkNDMtNDllNy1iYWY4LTY2MGZlZjkyZmQ3OCIsImRhdGEiOnt9LCJyYW5kb20iOiI5MGQ4NTE2YTE4OGViNDQ3YTFiZmMyYTFkZDlhZTZmMiJ9.CsJ0VRuQ90A1qzO1VJC1Q9eXFSp1N5UpeJlyr6otgXRPlf-I-jcwBJ6ytiBbbu8enCNJ2Ke6-oxNV8aeJ_AWIw

## Introduction

Modelling interactions between tropical forests and global climate, particularly the carbon cycle and other geophysical and geochemical processes, would greatly benefit from a more comprehensive description of vegetation dynamics from species level to community level across forest types. This is particularly true for tropical forests, which, despite their disproportionate role in the global carbon balance, are probably the least well described (Chen et al., 2020; Pan et al., 2024; Restrepo-Coupe et al., 2017). Key processes such as seasonal (phenological) variation in productivity (Wu et al., 2016), as well as the drivers and frequency of tree mortality events (Gora & Esquivel-Muelbert, 2021), are still insufficiently documented. To link biological processes occurring at tree level to broad-scale mechanistic models and spaceborne Earth observation methods, high-resolution spatial and temporal data are needed to capture individual tree dynamics over meaningful extents (Cushman et al., 2022).

Drone imagery is increasingly being used to study forests on the landscape scale, particularly through repeated acquisitions that enable temporal monitoring (Barbier et al., 2021; Park et al., 2019). The rapid advancement in the quality, affordability and user-friendliness of off-the-shelf drone equipment has made high-resolution aerial data more accessible than ever. Even without relying on high-end specialized sensors such as hyperspectral cameras or LiDAR, detailed RGB imagery can provide valuable insights into forest structure and dynamics (Cushman et al., 2022). However, the challenge lies in efficiently processing and transforming vast amounts of raw images into meaningful ecological and biophysical information. Indeed, even though individual photographs (e.g., collected by drones or any other means) can be geotagged (i.e., associated with a GPS position), they are only 2D projections of the 3D reality. Distortions due to viewing perspective and 3D object geometry make it difficult to precisely measure or localize objects. Using multiple views of a scene allows us to infer the 3D structure of objects thanks to the relative motion of near and far objects, and hence to project and eventually mosaic photographs. Stereophotogrammetry, or Structure from Motion (SfM) (Ulmann, 1979; Westoby et al., 2012), has become a standard tool for generating high-resolution topographic (3D) models and orthorectified imagery. While this technique is widely used in geosciences (Carrivick et al., 2016), its application in forestry is still emerging (Iglhaut et al., 2019). This process has been fully automated with modern computer vision algorithms and can now be performed efficiently on standard computing hardware, even with tens of thousands of high-resolution images. Earlier photogrammetric techniques required detailed information about camera parameters, such as lens properties (interior geometry) and position/orientation (exterior geometry). However, current software can autonomously detect keypoints and resolve camera acquisition (internal and external) geometry (incl. position, orientation, and lens properties) with minimal inputs.

To achieve high georeferencing and spectral precision, standard protocols currently necessitate: (i) the use of ground control points (GCPs) and/or differential GNSS positioning for centimeter-level absolute accuracy; and (ii) spectral calibration using reference targets or downwelling illumination sensors to convert raw spectral data into reflectance values. These protocols are widely implemented in industrial agriculture, where recent advances include miniaturized multispectral sensors with onboard illumination sensors and improved GNSS positioning using real-time kinematics (RTK). RTK enables real-time corrections through a reference network or an on-site base station, significantly enhancing positioning accuracy. Adapting these methods to tropical forests introduces specific challenges. Laying out spectral or spatial GCPs above the canopy is logistically challenging.

Reference GNSS networks are generally lacking, and dense canopy cover often obstructs RTK signal transmission. Furthermore, remote forested regions frequently lack internet access (4G/5G), further complicating real-time correction methods. Post-processing kinematic DNGSS (PPK) or subscription to global correction services (e.g., RTX) are options to reach centimetric positioning, but financial constraints in many tropical regions may necessitate cheaper solutions.

To address these challenges, we developed an automated processing workflow to derive useful quantitative information even in less favourable conditions, that is: without GCPs, DGNSS, spectral calibration, or multispectral sensors. The workflow combines existing freeware and proprietary software, and could eventually be completely open source pending some further adaptations. Our goal is to provide practical and accessible tools to detect and quantify changes in forest canopy at both individual tree and stand levels.

To track the phenology at the individual crown (or even branch) level, fine alignment of orthomosaics across dates is required. We used two recent methodologies, TIME-SIFT (Feurer & Vinatier, 2018) and Arosics (Scheffler et al., 2017) to improve the spatial coherence between image mosaics in the absence of DGNSS corrections. It is also essential to ensure spectral consistency across different sensors and over time to detect real biological changes rather than mere artifacts introduced by variations in equipment or processing methods. At a given spatial and spectral resolution, the spectral (and textural) signal is still impacted by environmental and technical factors, including cloud and tree shadows, atmospheric conditions, as well as sensor differences that modify the scene illumination and signal characteristics. To address spectral instability, an initial step involves adjusting color and white balance over overlap areas between aligned photos (ideally from multiples dates) to minimize the effects of varying lighting conditions. We also use vegetation indices (VIs), which are meant to be robust to illumination changes using ratios and proportions between spectral bands. We notably examine RGB-based indices that have increasingly been used to track phenological changes, such as the Green Chromatic Coordinate (GCC) (Lopes et al., 2016; Park et al., 2019).

We first test the processing chain on a series of drone acquisitions made in contrasted illumination conditions with both RGB cameras and multispectral sensors to quantify spectral and spatial instability and the effect of the implemented mitigation solutions. Second, we examine a longer time series of drone imagery to demonstrate the value of the spatial, spectral, and temporal information obtained at different scales. By integrating these methodological improvements, this research aims to improve the accuracy, affordability, and scalability of drone-based forest monitoring, ultimately contributing to a better understanding of the dynamics of tropical forests and their role in the global climate system.

## Materials and Methods

### Study Site

The study was carried out in a ∼50 ha area of interest surrounding the Guyaflux eddy flux tower at the Paracou permanent forest research facility in French Guiana (Bonal et al., 2008; Gourlet–Fleury et al., 2004). The site features a dense rainforest cover. The climate is classified as humid equatorial (rainfall c. 3000 mm/year), characterized by a distinct rainy season from December to July and a shorter primary dry season during the boreal summer, with a minimum monthly rainfall in September. Additionally, March typically experiences a brief reduction in rainfall. The soils are mostly nutrient-poor ferralitic soils.

The area of interest overlaps with 1617 ground-truthed tree crowns (Ball et al., 2023) and 10 0.5 ha square forest inventory plots (INRAe).

### Drone Imagery

303 photogrammetric flights were conducted using three different off-the-shelf DJI drones equipped with RGB and/or multispectral sensors. The missions were performed in autonomy using the UGCS flight planning software. In all cases, a target ground resolution of about 5 cm and an overlap of 80-90% were configured to allow stereophotogrammetric mosaicking.

The Phantom 4 Advanced (P4A) et Mavic 2 Pro (M2) drones comprise RGB stabilized cameras destined to general-purpose photography, with numerous manual and automated options adjusting exposition and white balance. Exposition was set to ‘Auto’ with an exposition value locked at 0 while white balance was left to vary (AWB). The images were saved in JPG format instead of RAW format, meaning that the white balance setting defined by the camera depending on the lighting conditions was definitely set in the images. This choice was made because not all systems allow saving in the RAW format and we thus had to choose the lowest common denominator.

The third drone was a Phantom 4 multispectral drone (P4M), which is more specifically designed for radiometric measurements. It has an RGB camera with less options (e.g. white balance cannot be modified), and 4 dedicated mono channel sensors covering broad spectral bands in the Green (550nm +/-40nm), Red (660nm +/-40nm), Red edge (735nm +/-10nm) and Near Infrared (790nm +/-40nm) regions. On its top, looking upward, the P4M also features an illumination sensor to capture incident radiation in the same five spectral bands, and hence to convert all acquired images to reflectance values (provided that the lighting conditions are homogeneous across the viewed scene).

Vegetation indices (VIs) were calculated as follows:

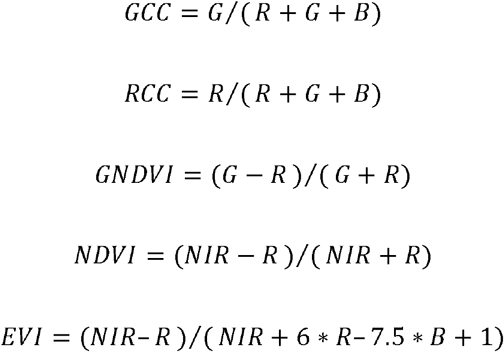

Where R, G, B, and NIR are digital numbers or reflectance values coming from the RGB camera or monochromatic sensors, depending on the case. The selection of indices is based on their broad applicability for vegetation monitoring, particularly in tropical vegetation phenology research, and their suitability for use with RGB sensors, which are commonly found in fixed cameras like Phenocams (Bradley et al., 2011; Lopes et al., 2016; Park et al., 2019). These indices are generally considered effective to i) stabilise the signal against modification in lighting and other acquisition conditions, and ii) highlight the biologically relevant information.

### Processing chain

We set up and automated a complete stereophotogrammetric workflow (Figure 1) based on existing software, covering all the way from multiple collections of RGB or multispectral images to the production of spatially and spectrally coherent multi-date orthomosaics, and the extraction of spectral variable and vegetation indices (VIs) at pixel, tree, or plot scale. The workflow is mainly coded in Python, but can be called from a simple R wrapper.

**Figure 1.**
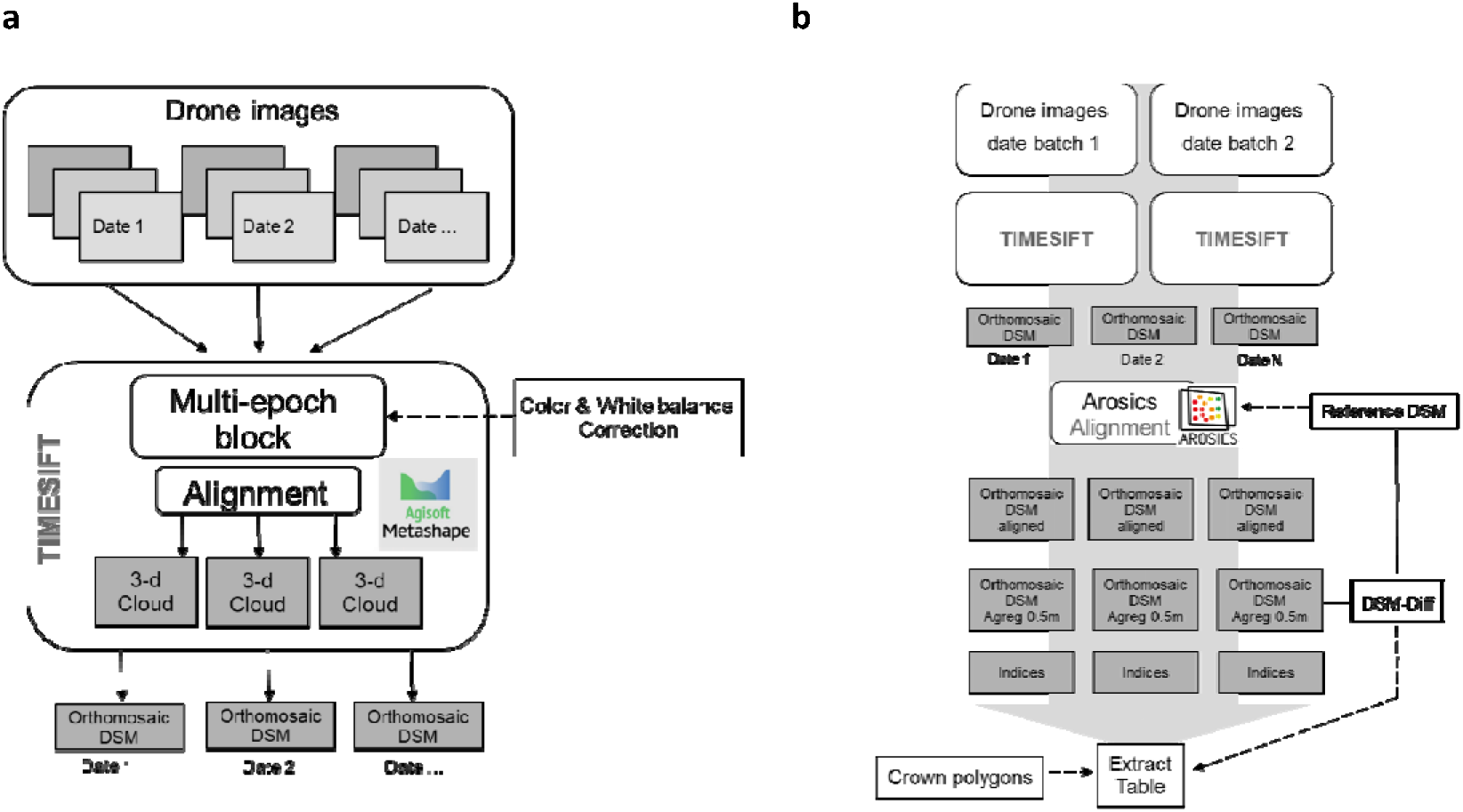
Diagram of the processing chain. a / details of the Time-SIFT multidate SfM approach. b/ residual spatial adjustments between Time-SIFT blocs and a reference (LiDAR) raster with Arosics and data extraction (case of ROIs defined by tree crowns polygons).

The core of the chain is Agisoft Metashape Pro, a widely used licensed software for automated stereophotogrammetric processing. The software is available both as a GUI and as a Python-based API. It features a rich library of functions and has become a reference for a broad user community. Using the principle of Structure from Motion (SfM), the software uses SIFT features to detect keypoints, reconstruct object geometry, and refine both the internal and external parameters of the cameras. Open-source alternatives exist, such as OpenDroneMap (Gbagir et al., 2023), Colmap (Maiwald et al., 2023) or MicMac (Brini et al., 2025; Filhol et al., 2019) and are also being adapted for our multi-date context (see Time-SIFT below).

#### Spatial Correction

As mentioned earlier, good-practice methods to ensure proper referencing of mosaics (differential GNSS and ground control points) may prove to be impractical or too expensive in the context of tropical forests. To ensure optimal spatial consistency between mosaics under DGNSS-denied conditions, two processing steps are implemented: Time-SIFT and Arosics. Time-SIFT is an adaptation of the normal SfM processing chain in which the initial image alignment phase is performed using photos from different dates (Feurer & Vinatier, 2018). The main postulate of this approach is that SIFT features are not only scale and rotation invariant, but also (to some extent) time invariant. Our workflow is heavily based on the Time-SIFT Python code released by the authors (https://doi.org/10.5281/zenodo.8359982)(Feurer & Vinatier, 2018). Two limitations may potentially limit the Time-SIFT approach: i) Although it ensures very high coherence between mosaics from a single processing block, absolute georeferencing still depends on the quality of the GNSS tags of the images. When differential GNSS is not used, the remaining shifts still generally require georeferencing the entire mosaic to a reference layer. ii) For long time series or large areas, computing limitations may prevent the alignment of all dates in a single Time-SIFT block. In these cases, it is useful to have a backup method that allows automated referencing of mosaics against a reference dataset. Arosics (Scheffler et al., 2017) is an open-source co-registration software based on raster data and Fourier cross-spectral analysis. It can be applied in two modes, global or local, depending on whether a single translation vector (orientation and magnitude) is computed for the whole image to reference, or whether local shift vectors are computed and then used in a polynomial transformation of the image to account for possible spatial distortions. In both cases, a window size needs to be set for the Fourier analysis, which was left here to the default value of 256×256 pixels. In the analysis presented here, an airborne LiDAR digital surface model (DSM) with 0.5m spatial resolution was used as a reference (acquired in 2019 by Altoa ltd.). Consequently, we chose to use stereophotogrammetric DSMs in Arosics to compute the global or local shifts. Remarkably, very similar results were obtained when the orthomosaics themselves. When using a local Arosics approach, the areas where spatial shifts are computed are generated following grid-like pattern with a specified granularity. Here we used a default grid resolution size of 1000 pixels (counted in the resolution of the reference layer, therefore 500 m). Both the resolution of the grid and the size of the window can be adjusted. The precision of resulting corrections is reported to be of 0.3 pixels (of the reference layer) or better (Scheffler et al., 2017).

To quantify the effect of Arosics and Time-SIFT spatial correction methods, individually or in combination, we first performed an internal validation using the vectors of local corrections suggested by Arosics at different steps to quantify the remaining shifts present between the mosaics and the reference LiDAR DSM. An independent external test was also conducted by mapping differences in surface height (DSM) between the produced mosaics and the reference LiDAR DSM. In such DSM difference maps, reconstruction anomalies and shifts are readily identifiable by visual inspection.

#### Color and illumination corrections

Spectral calibration using reference panels distributed over the area of interest is impractical when mapping a closed forest canopy. For RGB sensors, in addition to changes in illumination conditions, modifications in exposition values (iso, aperture, shutter speed) and white balance may also affect the recorded results. Agisoft proposes a color-calibration option that homogenises histograms (incl. white balance) between photos following the alignment phase. For MS data from the P4M drone, data from the multispectral illumination sensor placed on top of the drone were used in Agisoft to derive reflectance values.

For each flight, the average time and date of acquisition allowed to derive sun height. Although we could not distinguish between direct and diffuse illumination conditions, this variable was used to identify possible spectral biases induced by seasonal and interflight variation in sun height. We indeed expected that bidirectional reflectance (BRDF) effects would leave a signature in vegetation indices (Galvão et al., 2024).

### Test-sets

To test post-processing methods aiming at improving spatial and spectral coherence, we first selected small sets of acquisitions flights maximising the variability in illumination conditions (time of flight notably, but also atmospheric conditions and cloud shadows) for both the P4A and P4M platforms. Eighteen and seventeen flights were selected for each drone, respectively, with a balanced distribution in the morning, noon, and afternoon sessions. Full coverage of the 50-ha area of interest normally required three separate flights. Each acquisition flight comprises several hundred images, corresponding to about 20 minutes of acquisition time. Alignment of tests sets was performed as a single Time-SIFT block, pooling all dates for each sensor, or as date-wise blocks. Depending on the question addressed, mosaics were then generated per flight or per date. We also compared the quality of reconstructions obtained when pooling photos acquired in homogeneous vs. heterogeneous lighting conditions (time of day). SIFT features may indeed not remain invariant under very different illumination directions.

**Table 1.** Details of test-set flights from two different drones. P4A: DJI Phantom 4 Advanced. P4M Dji Phantom 4 Multispectral

| Date | Times | # Flights | Model | RGB | MS |
| --- | --- | --- | --- | --- | --- |
| 2020/10/23 | 9:28 - 10:49 | 3 | P4A | x |  |
| 2020/12/14 | 13:27 - 14:32 | 3 | P4A | x |  |
| 2020/12/15 | 14:31 - 15:32 | 3 | P4A | x |  |
| 2021/01/05 | 15:16 - 15:58 | 3 | P4A | x |  |
| 2021/03/03 | 14:04 - 15:58 | 4 | P4M | x | x |
| 2021/03/16 | 8:29 - 12:43 | 7 | P4M | x | x |
| 2021/04/06 | 9:15 - 12:53 | 6 | P4M | x | x |
| 2021/10/18 | 10:04 - 10:58 | 3 | P4A | x |  |
| 2021/11/23 | 10:32 - 11:23 | 3 | P4A | x |  |

### Long time series

To obtain a first glimpse of the signal obtained at crown and stand level over several seasons, we processed a time series of 84 dates (Appendix table) for the RGB sensors (301 flights in total) and 57 dates for the MS sensors (233 flights), spanning 3.5 years between October 2020 and May 2024. The dataset comprises 21 dates acquired with the P4A (RGB), 6 dates with the Mavic (RGB), and 57 dates with the P4M (RGB and MS). Time-SIFT blocks were made up of flights from random sets of four dates to maximize homogeneity throughout the time series. Mosaics were produced for each date, aggregating flights from the same day into a single mosaic.

### Data extraction

Although all mosaics were produced at a resolution of 5 cm per pixel, the analyzes in the following were performed on aggregated data at 0.5 m for faster processing. Depending on the analyses, pixel values were either extracted for a random set of 10,000 pixels or averaged on the scale of tree crowns or forest inventory plots.

### Analyses

Although the example datasets presented above were not specifically acquired for the purpose of a sensitivity analysis (e.g. with data from different sensors acquired simultaneously), our dataset is sufficient to document the relative portions of variance attributable to instrumental, environmental, and biological effects, as well as to spatial and spectral correction methods. Linearly regressing VIs values prior and after correction offers a way to assess if the data structure was globally affected.

## Results & discussions

### Spatial stability

Using a test set of 18 RGB acquisition flights spanning 6 dates (the P4A test set), we investigated sources of bias and the effect of methods to improve spatial consistency of the mosaics. We first based the analysis on the indication given by local correction shifts proposed by Arosics between the obtained stereophotogrammetric digital surface models (DSM) and the 2019 LiDAR reference DSM (0.5-m resolution). The spatial coherence obtained when aligning drone images date by date or when using a single Time-SIFT block for all flights were compared. The first approach both induced larger absolute and relative spatial errors (Figure 2a, blue boxes). Local shifts of up to 8 meters (i.e. 16 pixels of the reference LiDAR DSM) were even observed in an extreme case. A very high variance of local shift vectors was also observed in some mosaics (cf. the uncorrected 2021-10-18 mosaic). With the Time-SIFT approach (Figure 2b), global location errors were significantly reduced and homogenized (compare blue box plots of the figures on the left and right). Co-registration (via a global Arosics correction or the manual collection of a few reference points) remains necessary as long as differential GNSS corrections are not implemented, but as more GNSS positions are averaged across dates, the Time-SIFT approach effectively reduces global positioning bias. Thus, both relative and absolute spatial errors are attenuated.

**Figure 2.**
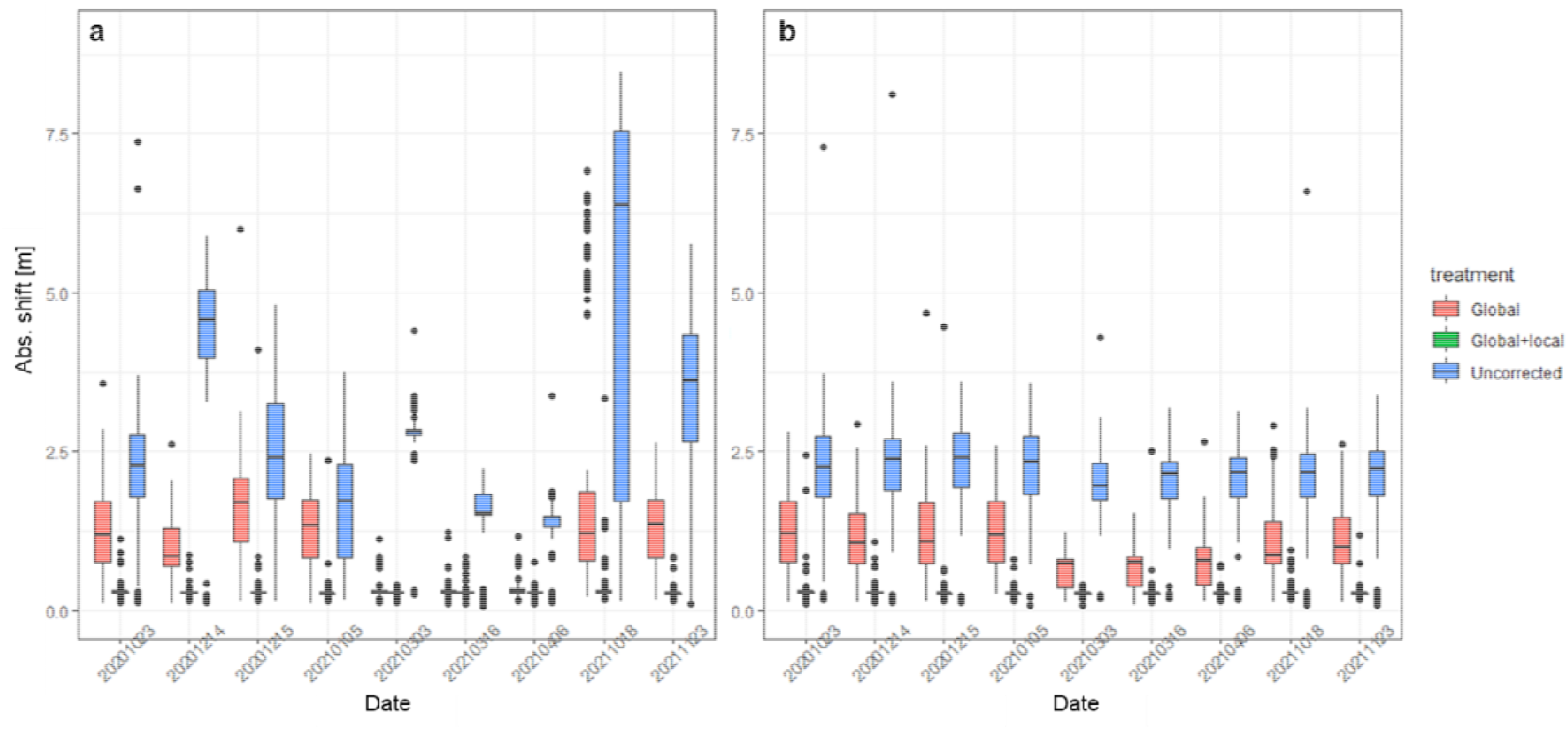
Measurement of the effect of misalignment mitigation methods (Arosics and Time-SIFT). Diagnostic provided by a box plot of absolute Arosics local shifts (computed every 500 m) against the reference 2019 ALS DSM (units in meters). a/ Date-wise mosaics (i.e. without Time-SIFT blocks). b Mosaics produced after Time-SIFT alignment. The treatments within each panel correspond to levels of Arosics processing, that is, without, after a global shift and after both global and local shifts.

Both Time-SIFT and local Arosics corrections proved effective in reducing the average value of local shift vectors (red and green bars low on the shift axis of both Figures 3a and b), as shown by a Kruskal-Wallis test (p-value < 2*10^−16^). Pairwise comparisons using the Wilcoxon rank sum test indicated that Time-SIFT and local Arosics corrections were both effective on their own and synergistically. However, the improvement from the combined use of both corrections (i.e. Time-SIFT + Local Arosics correction) was limited to an average reduction of the local shift vectors of a few centimeters.

**Figure 3.**
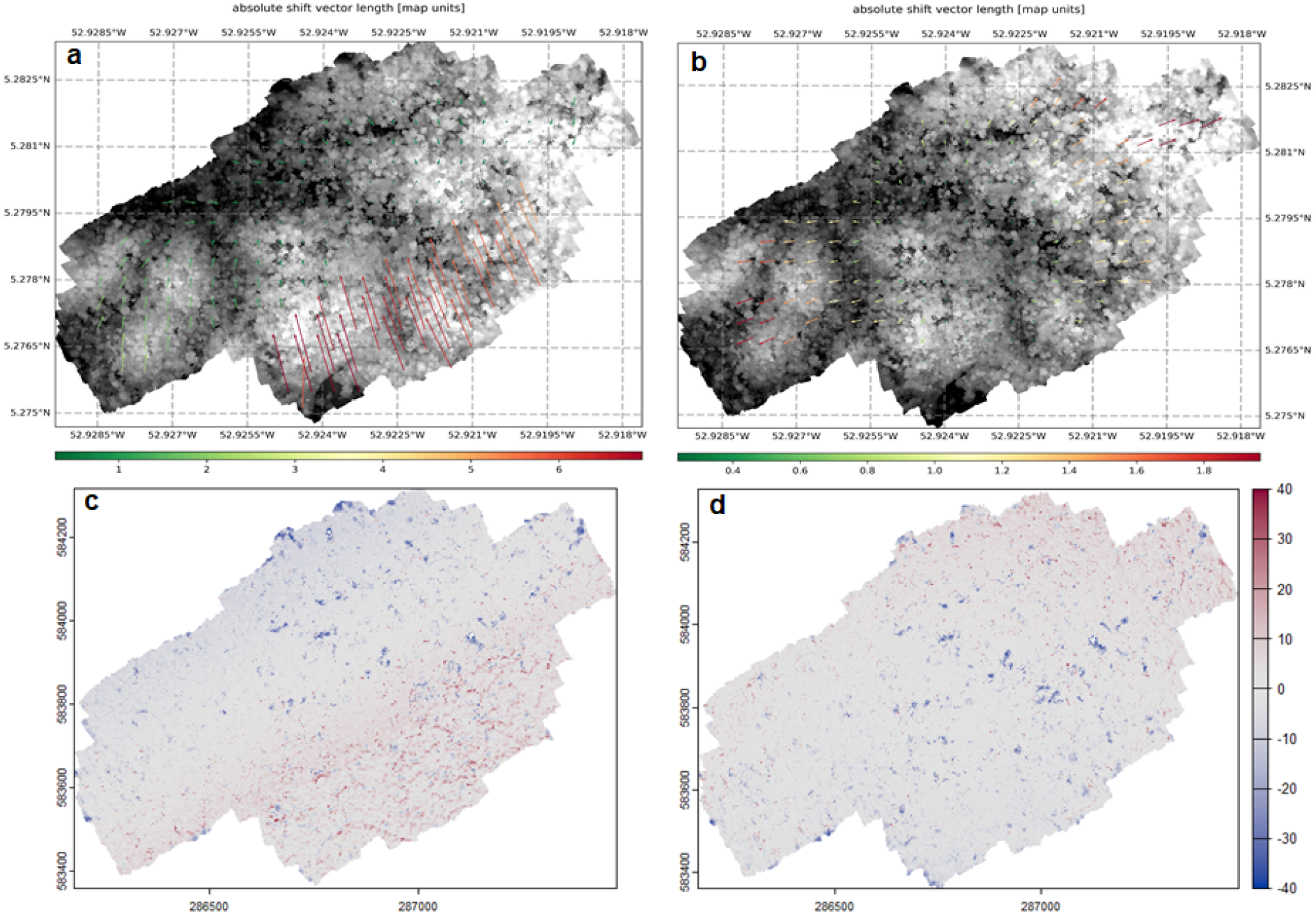
Spatial consistency of the P4M orthomosaic from 2021-10-18 after global shift depending on the reconstruction method (datewise or Time-SIFT). a & b/Arosics vector maps of local suggested corrections. The background layer is the reference ALS DSM in gray levels. The size of the vector representations was exaggerated (factor 10) for visibility. Notice the change in color scale between vector maps. The color scale is in meters. Spatial reference in geographic coordinates (WGS84). a/ Datewise mosaic (i.e. without Time-SIFT group). b/ Mosaic from Time-SIFT group. Notice the large suggested correction vectors in the south part. c & d/ DSM difference maps. Color scale in meters of vertical difference. Spatial reference in UTM 33N. c/ Datewise mosaic (i.e. without Time-SIFT group). Notice the systematic positive deviations (red) in the south part. d/ mosaic from the Time-SIFT group. The large negative deviations (blue) correspond to tree and branch fall since 2019.

A reason for using Time-SIFT and not just a Local Arosics for each single date mosaics is that by increasing data redundancy and reducing some holes in the photo grid, Time-SIFT blocks allow to mitigate occasional (but potentially large) reconstruction anomalies. For example, the mosaic of 2021-10-18 had the largest local deviations (Figure 2a). The date-wise mosaic (produced without the Time-SIFT approach) showed a large inconsistency between the northern and southern parts of the mosaic (Figure 4). The local corrections suggested by Arosics (after a global correction, Figure 3a) relative to the ALS reference map of 2019, could reach values of more than 6 meters in the southern part of the mosaic. Similarly, the DSM difference map (Figure 3c) detects many positive vertical anomalies in this area, a typical feature that indicates a spatial change between the two compared DSMs. On the other hand, the mosaic produced for the same date out of a Time-SIFT group (Figure 3b & d) shows much reduced local deviations in both maps (small shift vectors and limited positive DSM deviations). Interestingly, for the single-date mosaic, even after a local Arosics correction (Figure 4), some DSM deviations remained, especially at the interface between two different flights. This highlights the importance of a good initial 3D reconstruction in Agisoft, and the role the Time-SIFT approach can have in this regard. The local Arosics correction would need to be applied very locally (i.e. on a very tight grid) to compensate for a poor initial reconstruction. We do not show the local Arosics vectors here, as they are all very small, since the correction has already been applied.

**Figure 4.**
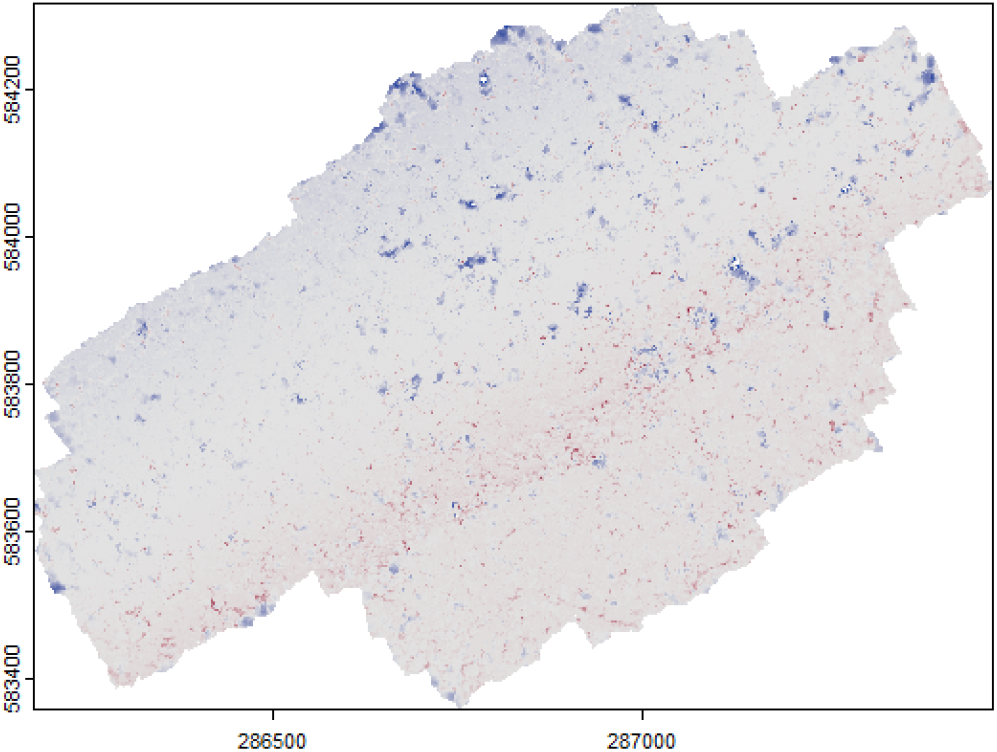
Spatial consistency (DSM difference map) after local shift correction in Arosics for the P4A 2021-10-18 datewise orthomosaic (i.e., without Time-SIFT group). For reference see Figure 3. Many positive deviations (red) are still present.

Because SfM is based on the detection of invariant features in the images, strong illumination changes can impact the quality of reconstructions. To explore how changes in lighting conditions can affect the accuracy of spatial alignment, we built Time-SIFT blocks that combine flights acquired at specific times in the day. When using only morning or only afternoon flights, we observed a very significant reduction (p-value <10^−4^, Kruskal-Wallis and Pairwise Wilcoxon rank sum tests) in the proposed absolute shifts in local Arosics corrections, with average absolute shift reductions of up to 30 cm. We therefore recommend limiting acquisition to a few (constant) hours of the day when it is feasible and when a particularly high spatial coherence is required through time. Contrary to (Iglhaut et al., 2019), we recommend avoiding mid-day (hotspot) conditions, which alter both the spectral properties and the detectability of the tree crown. Aiming at constant atmospheric (overcast or sunny) conditions for many repeated flights over long monitoring periods is obviously unrealistic.

### Spectral stability/sensitivity

We analysed the spectral stability and the effect of spectral mitigation methods for mosaic data aggregated at either pixel, crown, or plot levels. From here on, the spatial corrections (Time-SIFT and local Arosics) discussed above are considered to be applied and optimal.

### RGB sensitivity

#### Inter-flight variance and color calibration

The application of the color-calibration procedure available in Agisoft on a mosaic of three different flights from the P4A test-set is illustrated in Figure 5. The effect of color calibration is apparent in the RGB mosaic, but even more so in the GCC map, where the boundaries between the three flights are largely eliminated in the corrected version.

**Figure 5.**
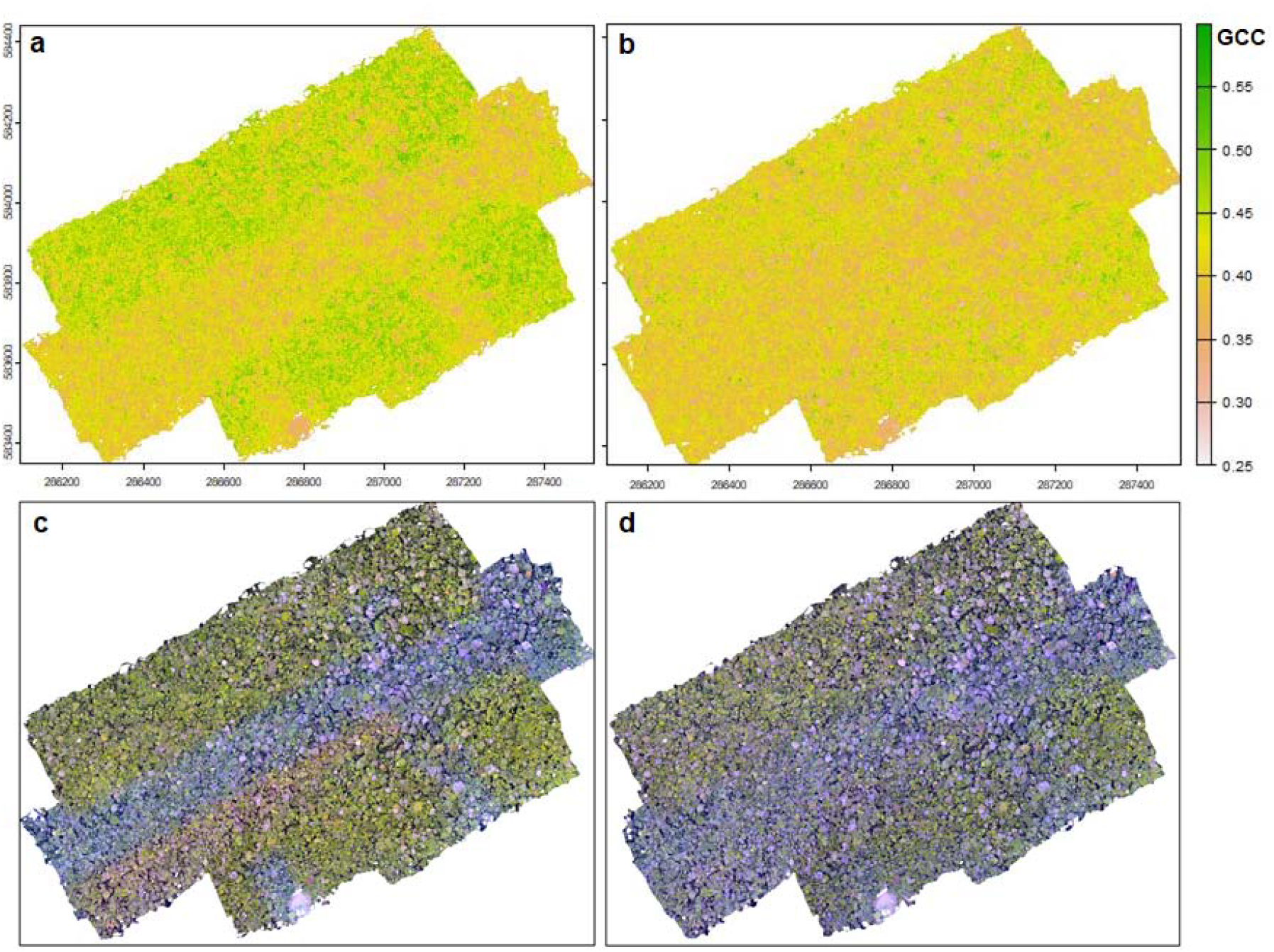
GCC index and RGB color composite from RGB Mosaic of three flights (2020-11-05), a-c/ without color calibration, b-d/ with color calibration

**Figure 6.**
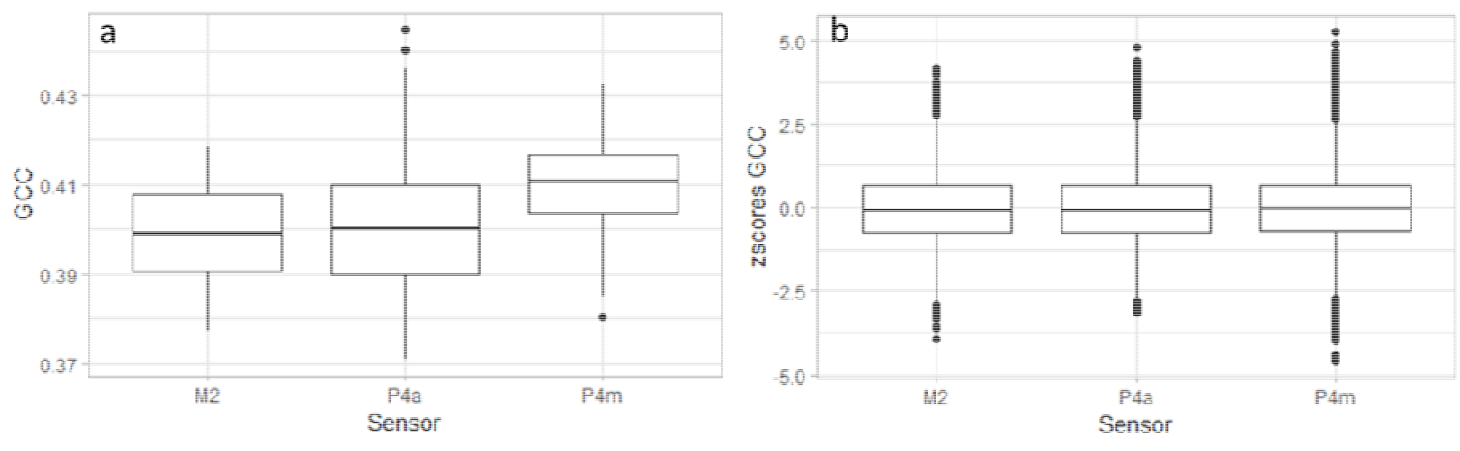
Box plot illustrating the effect of the sensor on the GCC index. a/ without intervention. All pairwise differences are very significant according to Tuckey HSD (alpha < 10^−6^). Anova on sensor effect, R2=0.13. b/ after standardisation.

To further evaluate the benefits of this spectral correction, we compared the effect of color and white balance corrections across flights and dates (using the Time SIFT block) to data obtained from independent mosaics produced per flight (hence without correction). Applying the color calibration to multiple dates and flights yielded a ∼50% reduction in the variance of the values of digital numbers (DN) and the derived indices. For example, for the GCC index, the mean sum of squares (MSS) of the ‘flight effect’ went from 3.37 to 1.54. For the RCC index, the MSS went from 20.47 to 10.2. Note that beyond differences between flights, color- and white-balance correction also have the potential to decrease spectral variability within each flight (e.g. due to cloud shadows).

However, the overall structure of the signal remained relatively stable, with a linear regression between corrected GCC and raw GCC producing a R2 = 0.71 (y=0.08+0.8x) and between corrected RCC and raw RCC a R2=0.74 (0.06+0.76x). Therefore, reducing noise sources using intercalibration with the color correction does not eliminate potentially informative patterns present in the data, as will be later illustrated with the long time series.

#### Sensor effect

Using the long RGB time series, we detected a very significant sensor effect on the distribution of VI values, even after color calibration. The latter appears to be applied separately to the different sensors recognized by Agisoft. An ANOVA on sensor effect proved very significant (p-value <10^−6^) on all VIs, with R2 values between 0.12 and 0.54. Both the variance and the mean of the VI value distributions changed between the sensors. A pairwise Tuckey HSD test showed significant differences between all sensors. A simple standardisation by sensor effectively removed the effect. In practice, using such drastic correction implies having time series of sufficient length and representativity for each sensor to avoid removing relevant biological information. In our case, this could be problematic for the shorter Mavic 2 time series. A better approach would be to use precomputed offset/gains, for instance, using intercalibration flights acquired nearly synchronously between sensors.

#### Sun height effect

Despite a broad range of average sun height conditions spanning from 20 to 90 ° depending on the flights, we were unable to identify clear trends in the average values for most studied vegetation indices (GCC, NDVI, GNDVI, RCC) that would be induced by sun height variation, either in the corrected or uncorrected test sets. However, a significant effect of sun height was found on EVI (R2 = 0.3, df = 13, p-value = 0.03) for both uncorrected and illumination corrected data. These results confirm satellite-derived observation that EVI was more sensitive to directional effects than other indices (Galvão et al., 2024; Kattenborn et al., 2024). The long time series furthermore allowed to show that the temporal sampling of sun height was haphazard (Figure 7), and therefore unlikely to produce a structured bias induced by changing sun height across seasons, which is a known issue for sun synchronous satellite data (Morton et al., 2014).

**Figure 7.**
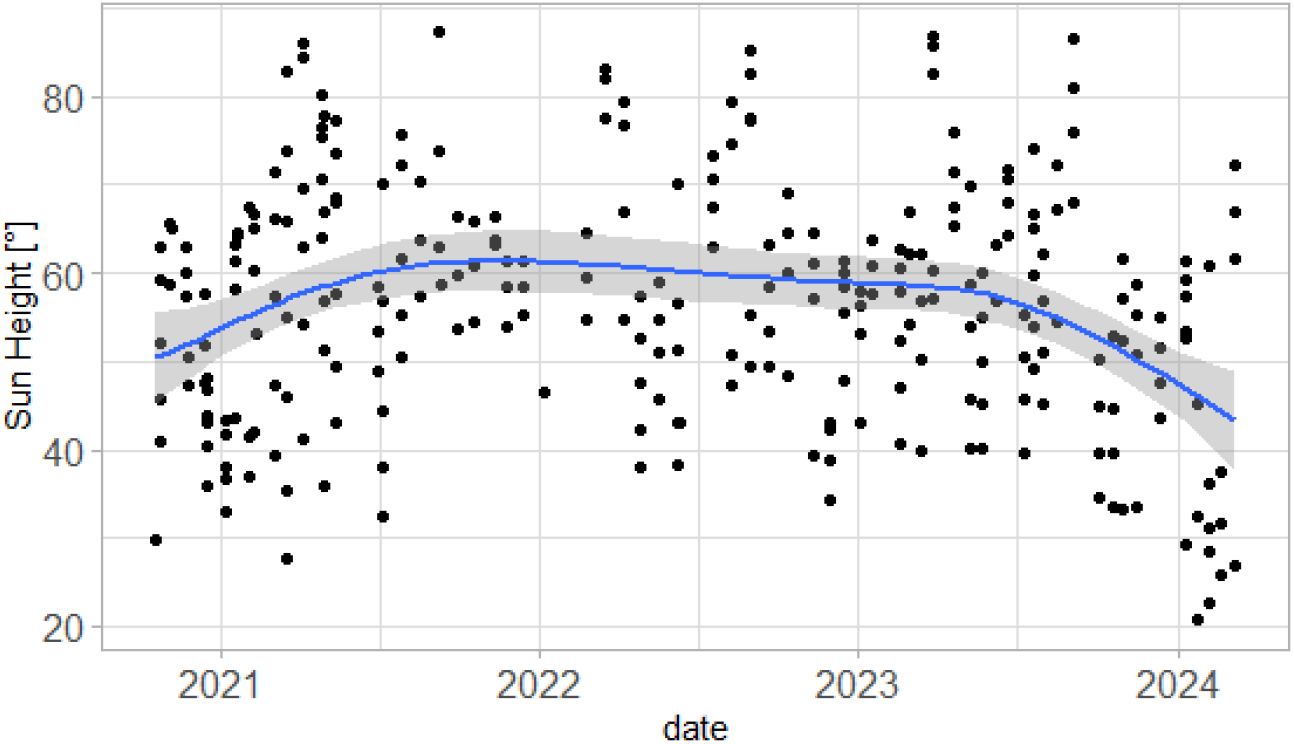
Temporal sampling of Sun Height angle. Fit line obtained with general additive modelling.

### Sensitivity of multispectral data

For VIs derived from the P4M multispectral sensors, exposition values and white balance are fixed and calibrated for radiance measurements. Consequently, color calibration resulted in a smaller reduction in interflight variance compared to other sensors, between 19 and 28.4% for GCC, between 27 and 42% for NDVI, and even an 8.5 to 15% increase in variance for RCC. However, when using the illumination sensor to further correct interdate changes in atmospheric and illumination conditions, the noise reduction proved consequent, with a 73.3 to 79.4% reduction for GCC, 86.3% to 89.7% reduction for NDVI, and 27.7 to 49 % reduction for RCC.

Given the magnitude of the above corrections, we tested how much the data structure was impacted, by regressing corrected to noncorrected VI values. For example, for GCC after color calibration, the relationship between corrected and uncorrected values followed y=0.06+0.81x with R2=0.7. However, with the calculation of reflectance values using the illumination sensor, the correlation dropped (y=0.32+0.57x, R2=0.34), showing a significant change in the structure of the data. The results were comparable for the RCC and NDVI indices.

To have a clearer understanding of the correlation structure between VIs and processing levels, we compared the indices obtained using the RGB sensor (after color calibration) and the MS monochromatic sensors after calibration with the illumination sensor. Both types of data are indeed acquired simultaneously by the P4M drone. In the case of GCC, the correlation was quite good (y=-0.014+1.32x R2=0.73). However, for RCC, it was very low (y= 0.05+0.42x, R2=0.07). We extracted VIs for each crown per date and per sensor (RGB or MS) by averaging the pixel values within the delineated crowns. The correlation matrix between the different VIs derived from color-corrected RGB sensor data or illumination corrected MS data (Figure 8a) shows that overall, the various VIs indicative of greenness or chlorophyll concentration (GCC, NDVI, GNDVI) are intercorrelated, with the notable exception of EVI. The EVI appears to display an important independent component of variability (along the third axis of a PCA, Figure 8c), which is to be treated with some caution considering its directional sensitivity. The lack of correlation between MS-RCC and RGB-RCC is also clearly visible (Figure 8b). Notice the strong negative correlation between MS-RCC and the greenness indices, which is consistent with expectation. The low consistency for RCC between sensors and processing levels is likely a result of the lower signal of vegetation in the red (due to high absorption) and should be kept in mind when using this information.

**Figure 8.**
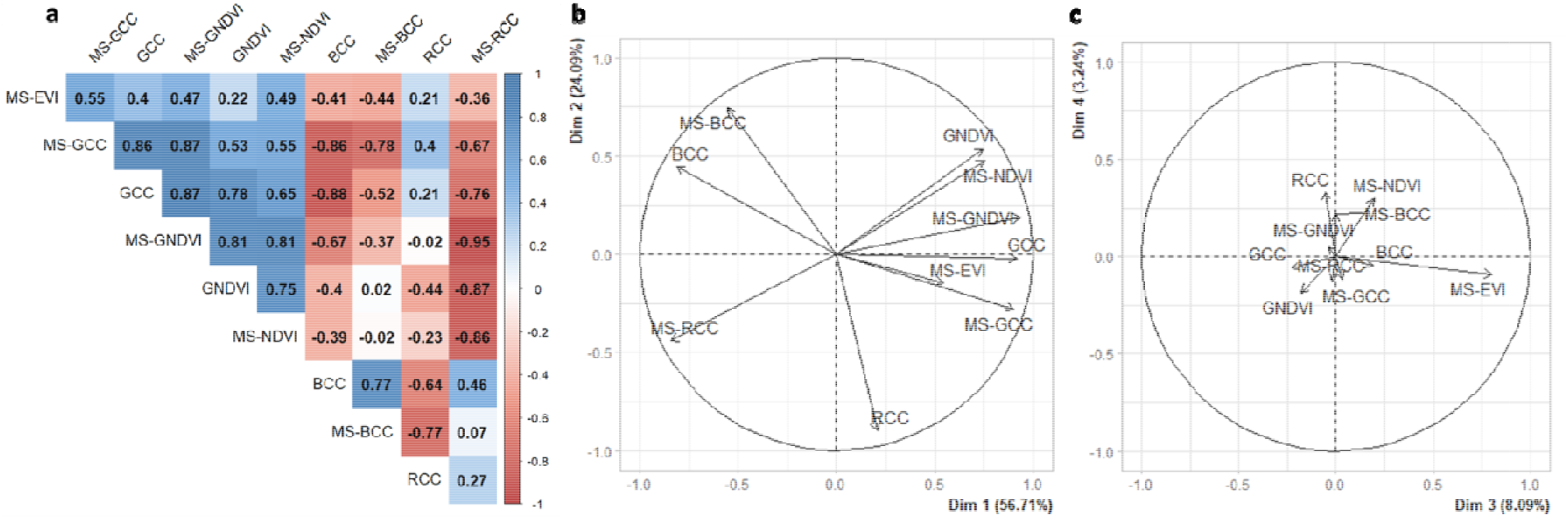
Variable correlations in the first plane of a principal component analysis performed on crown averaged vegetation indices obtained from the RGB and MS sensors of the P4M over all available dates (N=412 371 observations).

On the other hand, indices based on the green and NIR wavelengths appear to carry a consistent signal across sensors. A major conclusion is that cheaper RGB sensors produce usable data for GCC, at least in terms of relative changes, and that the post-processing (color calibration) proposed by Agisoft does help improving the signal.

A major advantage and purpose of MS sensors is to provide (absolute) calibrated reflectance values that are in theory comparable to satellite products. See however Cottrell et al. (2024) for some limitation of existing high-end devices. Moreover, using the information content of spectral bands that are thematically important but for which the signal is relatively weaker (e.g. the red band and derived indices) may require using MS sensors with proper incident light correction.

Importantly, drone surveys contain much more than just vegetation index (VI) values, offering high spatial (3D) and temporal resolution for enhanced data insights. Phenophases can be interpreted visually or automatically using textural and temporal context information. Furthermore, we have shown here that both spatial and spectral imperfections can in large part be mitigated at the postprocessing stage, thanks to the automated processing chain.

### Illustration of time series at the stand and tree levels

We illustrate the signal obtained using the long time series of RGB and MS sensors. We focus on the GCC index. At stand level (Figure 9) we can clearly observe an annual cyclical variation peaking in the dry season (August-Sept.-Oct), highlighted by a fit from a general additive model of the data over time. Some discrepancies are observed between the RGB (light green curve) and MS-derived versions of the VI. The level of inter-date variability does not seem to be much lower in the multispectral data compared to the RGB data.

**Figure 9.**
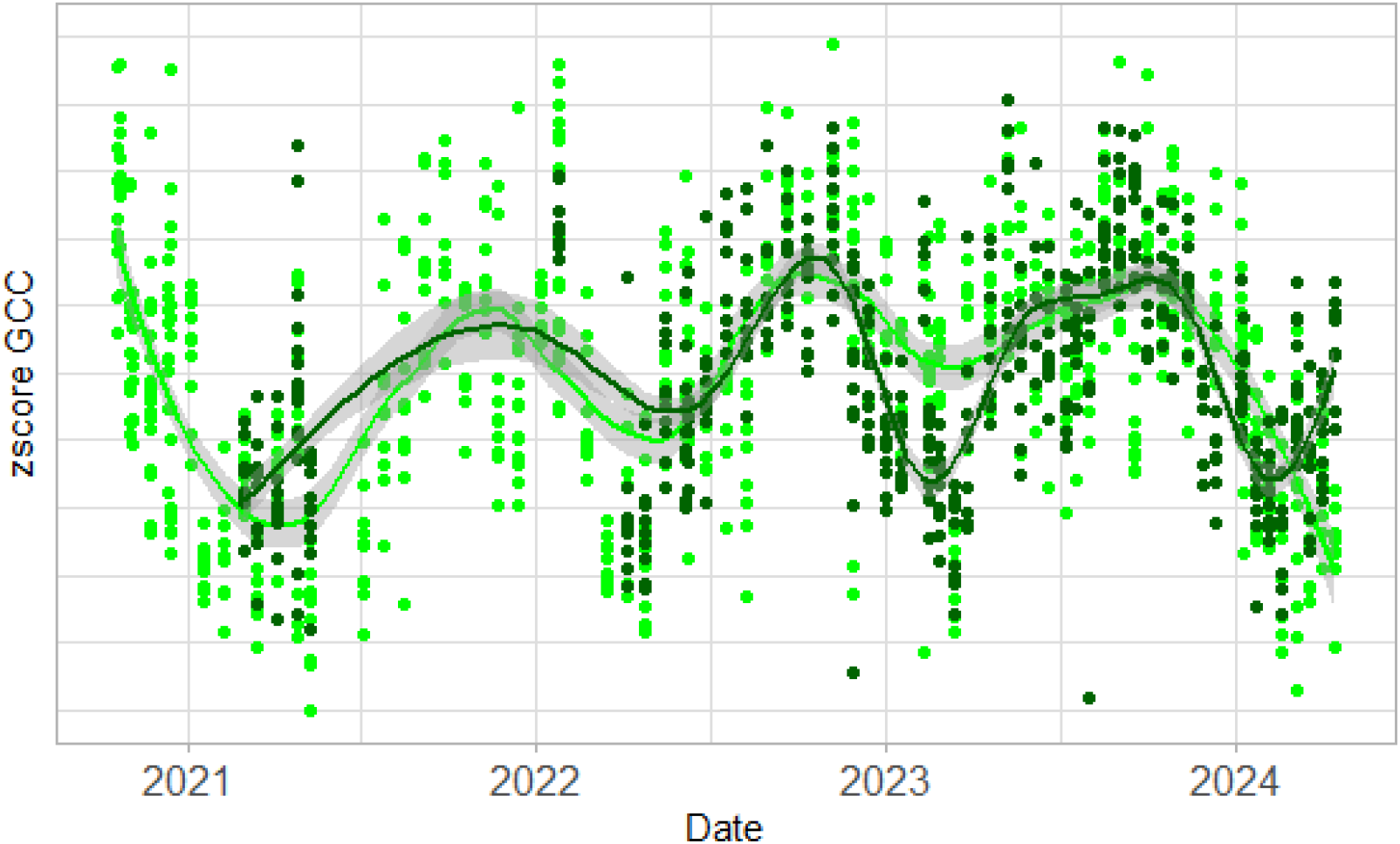
Long time series of plot level greenness (GCC z scores) over 8 plots of 80×80 m (INRA plots) distributed eastward of the Paracou Guyaflux tower. Light green: RGB derived GCC values after color-calibration and sensor-wise standardisation. Dark green: illumination-corrected GCC (and standardized) values from MS sensors. Fit line obtained with General Additive Modelling.

We also represent visual variations at tree level for the Parkia nitita species (Figure 10), showing gradual or brutal changes in foliage color and apparent density, with phases of defoliation and flushes. We can notice a desynchronised branch as well that defoliates and flushes slightly later than the crown. Figure 11 illustrates the synchronized and yearly greenness pattern observed for the local population of Recordoxylon speciosum tree species. This (arbitrarily selected) species shows a clear defoliation/refoliation pattern followed by a gradual decrease in greenness.

**Figure 10.**
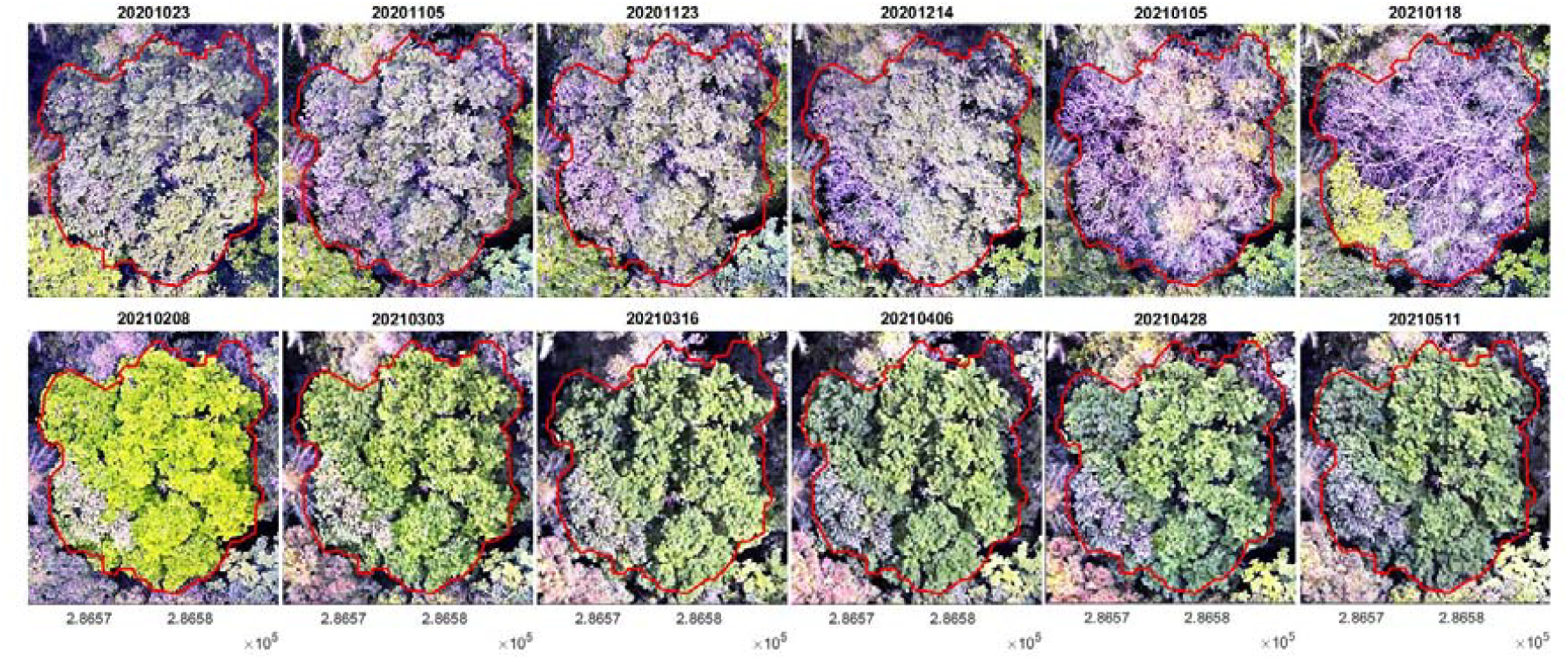
Visuals at tree level of a Parkia nitida tree, showing changes in foliage color and density, as well as desynchrony in vegetative phenology. Note that an additional histogram stretch was applied for each extract for visualisation.

**Figure 11.**
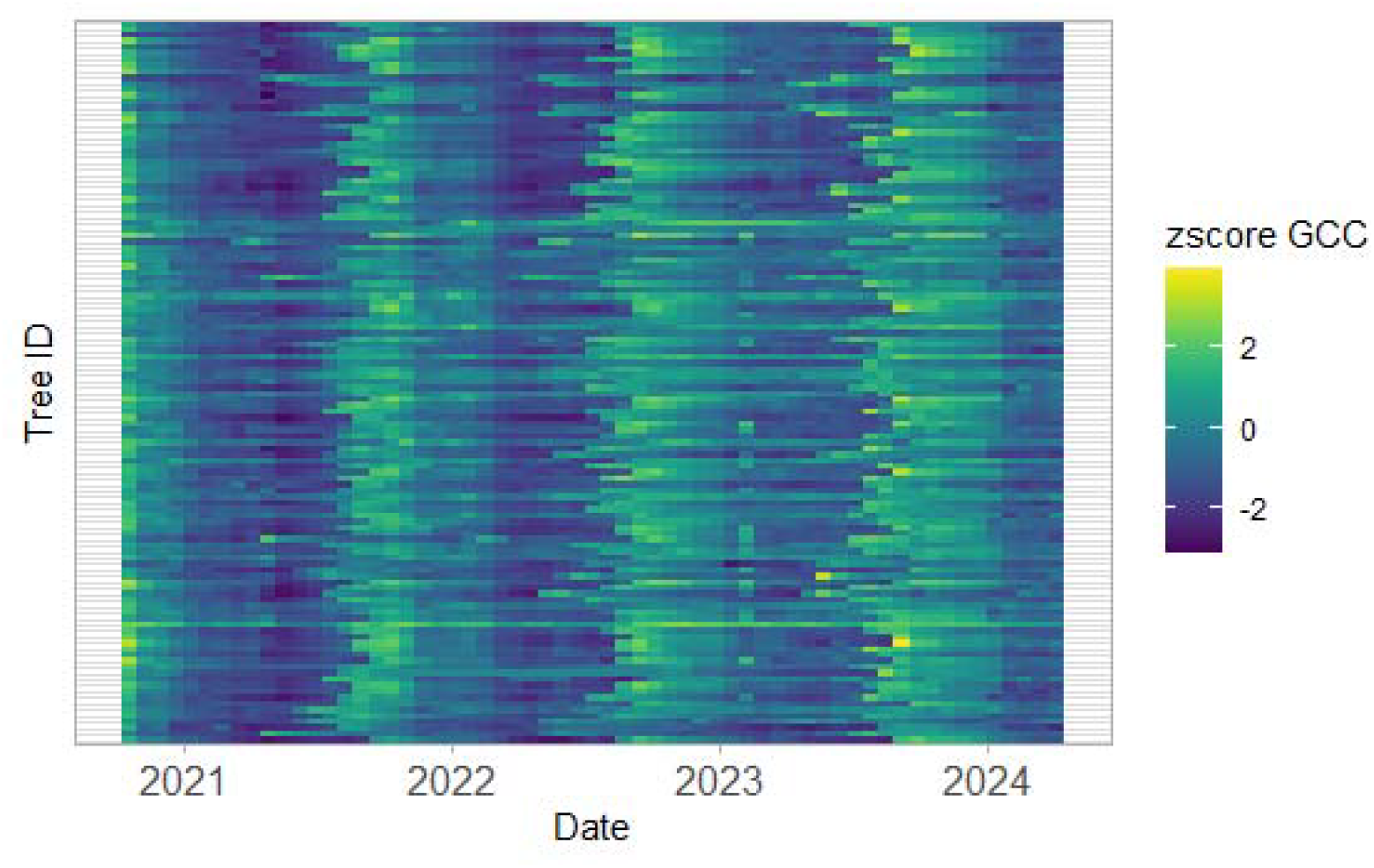
Heat map of tree-level GCC values (each line is a single digitized crown of Recordoxylon speciosum) through time (long time series from RGB cameras).

These illustrations demonstrate the existence of quantitative variation in vegetation spectral and textural properties, perceptible at both tree level and stand level. The spatial and spectral stability of the obtained information opens the way to numerous applications in biology and ecology studies at a range of study scales.

## Conclusions

We established the feasibility and interest of using light drones (incl. with a simple RGB camera and standard GNSS) for tree and stand-level monitoring in the particularly challenging context of tropical forests. Qualitative and quantitative information on phenology can be collected using spectral bands and derived spectral indices for which the vegetation signal is strong. Indeed, both the spectral and spatial precision allowed by the processing chain are sufficient to evidence subtle greenness changes that can be related to changes in leaf area or age (Wu et al., 2016). Beyond the use of the sole greenness signal, height variations as well as visual interpretation and computer vision algorithms will allow one to extract a wealth of information regarding the timing and patterns of tree primary growth, vegetative and (for certain species) reproductive phenology, branch fall, crown expansion, and tree death.

We provide a complete end-to-end processing chain that also includes tools for the visualisation, interpretation, and statistical analysis of the extracted data.

## Acknowledgements

This study benefited from an “Investissement d’Avenir” grant managed by the Agence Nationale de la Recherche (CEBA, ref. ANR-10-LABX-25-01), via project PHENOBS. We also acknowledge the support of the UE Biodiversa+ BiodivMon program (Project Coforfunc). We are thankful to Ilona Clocher, Jean-Louis Smock, Jean-Yves Goret, Florian Jeanne and Julien Engel for their help with drone data acquisition and/or processing. Access to the Paracou site and infrastructure (https://paracou.cirad.fr) was granted by CIRAD/Ecofog, and we thank Géraldine Derroire and all Phenobs project participants. We are grateful to Raphaël Pélissier for helping in the initial shaping of the Phenobs project and for his continued support.

## References

Ball, J. G. C., Hickman, S. H. M., Jackson, T. D., Koay, X. J., Hirst, J., Jay, W., Archer, M., Aubry-Kientz, M., Vincent, G., & Coomes, D. A. (2023). Accurate delineation of individual tree crowns in tropical forests from aerial RGB imagery using Mask R-CNN. Remote Sensing in Ecology and Conservation, n/a(n/a). 10.1002/rse2.332

Barbier, N., Ball, J., Clocher, I., Poilvé, H., Verley, P., & Vincent, G. (2021). Sensing Tropical Forest Phenology and Productivity from the Field to the Satellite. 2021 IEEE International Geoscience and Remote Sensing Symposium IGARSS, 716–719. 10.1109/IGARSS47720.2021.9554372

Bonal, D., Bosc, A., Ponton, S., Goret, J.-Y., Burban, B., Gross, P., Bonnefond, J.-M., Elbers, J. A. N., Longdoz, B., & Epron, D. (2008). Impact of severe dry season on net ecosystem exchange in the Neotropical rainforest of French Guiana. Global Change Biology, 14(8), 1917–1933.

Bradley, A. V., Gerard, F. F., Barbier, N., Weedon, G. P., Anderson, L. O., Huntingford, C., Aragao, L. E. O. C., Zalazowski, P., & Arai, E. (2011). Relationships between phenology, radiation and precipitation in the Amazon region. Global Change Biology, 17(6), 2245–2260.

Brini, I., Feurer, D., Tebourbi, R., Vinatier, F., & Abdelfattah, R. (2025). Evaluating the potential of the Time-SIFT approach using Pléiades satellite imagery for 3D change detection. Journal of Applied Remote Sensing, 19(2), 024501–024501.

Carrivick, J. L., Smith, M. W., & Quincey, D. J. (2016). Structure from Motion in the Geosciences. John Wiley & Sons. https://books.google.com/books?hl=fr&lr=&id=tZCvDAAAQBAJ&oi=fnd&pg=PR3&dq=Structure+from+Motion+in+the+Geosciences&ots=EjNiYQa_2c&sig=eI6fBO4Ccg74yxCdsn_0TosChIU

Chen, X., Maignan, F., Viovy, N., Bastos, A., Goll, D., Wu, J., Liu, L., Yue, C., Peng, S., & Yuan, W. (2020). Novel representation of leaf phenology improves simulation of Amazonian evergreen forest photosynthesis in a land surface model. Journal of Advances in Modeling Earth Systems, 12(1), e2018MS001565.

Cottrell, B., Kalacska, M., Arroyo-Mora, J.-P., Lucanus, O., Inamdar, D., Løke, T., & Soffer, R. J. (2024). Limitations of a Multispectral UAV Sensor for Satellite Validation and Mapping Complex Vegetation. Remote Sensing, 16(13), 2463.

Cushman, K. C., Detto, M., García, M., & Muller-Landau, H. C. (2022). Soils and topography control natural disturbance rates and thereby forest structure in a lowland tropical landscape. Ecology Letters, 25(5), 1126–1138. 10.1111/ele.13978

Feurer, D., & Vinatier, F. (2018). Joining multi-epoch archival aerial images in a single SfM block allows 3-D change detection with almost exclusively image information. ISPRS Journal of Photogrammetry and Remote Sensing, 146, 495–506.

Filhol, S., Perret, A., Girod, L., Sutter, G., Schuler, T. V., & Burkhart, J. F. (2019). Time-Lapse Photogrammetry of Distributed Snow Depth During Snowmelt. Water Resources Research, 55(9), 7916–7926. 10.1029/2018WR024530

Galvão, L. S., Arlanche Petri, C., & Dalagnol, R. (2024). Coupled effects of solar illumination and phenology on vegetation index determination: An analysis over the Amazonian forests using the SuperDove satellite constellation. GIScience & Remote Sensing, 61(1), 2290354. 10.1080/15481603.2023.2290354

Gbagir, A.-M. G., Ek, K., & Colpaert, A. (2023). OpenDroneMap: Multi-platform performance analysis. Geographies, 3(3), 446–458.

Gora, E. M., & Esquivel-Muelbert, A. (2021). Implications of size-dependent tree mortality for tropical forest carbon dynamics. Nature Plants, 7(4), 384–391. 10.1038/s41477-021-00879-0

Gourlet–Fleury, S., Ferry, B., Molino, J. –F., Petronelli, P., & Schmitt, L. (2004). Experimental plots: Key features. In Ecology and management of a Neotropical rainforest: Lessons drawn from Paracou, a long-term experimental research site in French Guiana (pp. 1–60). Elsevier.

Iglhaut, J., Cabo, C., Puliti, S., Piermattei, L., O’Connor, J., & Rosette, J. (2019). Structure from Motion Photogrammetry in Forestry: A Review. Current Forestry Reports, 5(3), 155–168. 10.1007/s40725-019-00094-3

Kattenborn, T., Wieneke, S., Montero, D., Mahecha, M. D., Richter, R., Guimarães-Steinicke, C., Wirth, C., Ferlian, O., Feilhauer, H., Sachsenmaier, L., Eisenhauer, N., & Dechant, B. (2024). Temporal dynamics in vertical leaf angles can confound vegetation indices widely used in Earth observations. Communications Earth & Environment, 5(1), 1–11. 10.1038/s43247-024-01712-0

Lopes, A. P., Nelson, B. W., Wu, J., de Alencastro Graça, P. M. L., Tavares, J. V., Prohaska, N., Martins, G. A., & Saleska, S. R. (2016). Leaf flush drives dry season green-up of the Central Amazon. Remote Sensing of Environment, 182, 90–98.

Maiwald, F., Feurer, D., & Eltner, A. (2023). Solving photogrammetric cold cases using AI-based image matching: New potential for monitoring the past with historical aerial images. ISPRS Journal of Photogrammetry and Remote Sensing, 206, 184–200.

Morton, D. C., Nagol, J., Carabajal, C. C., Rosette, J., Palace, M., Cook, B. D., Vermote, E. F., Harding, D. J., & North, P. R. (2014). Amazon forests maintain consistent canopy structure and greenness during the dry season. Nature. http://www.nature.com/nature/journal/vaop/ncurrent/full/nature13006.html

Pan, Y., Birdsey, R. A., Phillips, O. L., Houghton, R. A., Fang, J., Kauppi, P. E., Keith, H., Kurz, W. A., Ito, A., Lewis, S. L., Nabuurs, G.-J., Shvidenko, A., Hashimoto, S., Lerink, B., Schepaschenko, D., Castanho, A., & Murdiyarso, D. (2024). The enduring world forest carbon sink. Nature, 631(8021), 563–569. 10.1038/s41586-024-07602-x

Park, J. Y., Muller-Landau, H. C., Lichstein, J. W., Rifai, S. W., Dandois, J. P., & Bohlman, S. A. (2019). Quantifying Leaf Phenology of Individual Trees and Species in a Tropical Forest Using Unmanned Aerial Vehicle (UAV) Images. Remote Sensing, 11(13), 1534. 10.3390/rs11131534

Restrepo-Coupe, N., Levine, N. M., Christoffersen, B. O., Albert, L. P., Wu, J., Costa, M. H., Galbraith, D., Imbuzeiro, H., Martins, G., & da Araujo, A. C. (2017). Do dynamic global vegetation models capture the seasonality of carbon fluxes in the Amazon basin? A data-model intercomparison. Global Change Biology, 23(1), 191–208.

Scheffler, D., Hollstein, A., Diedrich, H., Segl, K., & Hostert, P. (2017). AROSICS: An automated and robust open-source image co-registration software for multi-sensor satellite data. Remote Sensing, 9(7), 676.

Ulmann, S. (1979). The interpretation of structure from motion. Proceedings of the Royal Society of London. Series B. Biological Sciences, 203(1153), 405–426. 10.1098/rspb.1979.0006

Westoby, M. J., Brasington, J., Glasser, N. F., Hambrey, M. J., & Reynolds, J. M. (2012). ‘Structure-from-Motion’ photogrammetry: A low-cost, effective tool for geoscience applications. Geomorphology, 179, 300–314. 10.1016/j.geomorph.2012.08.021

Wu, J., Albert, L. P., Lopes, A. P., Restrepo-Coupe, N., Hayek, M., Wiedemann, K. T., Guan, K., Stark, S. C., Christoffersen, B., Prohaska, N., & others. (2016). Leaf development and demography explain photosynthetic seasonality in Amazon evergreen forests. Science, 351(6276), 972– 976.

